# Human primary visual cortex shows larger population receptive fields for binocular disparity-defined stimuli

**DOI:** 10.1101/2021.03.10.434843

**Authors:** Ivan Alvarez, Samuel A. Hurley, Andrew J. Parker, Holly Bridge

**Author notes:** Corresponding author Holly Bridge, Wellcome Centre for Integrative Neuroimaging, FMRIB Centre, John Radcliffe Hospital, Oxford, UK, OX3 9DU.

## Abstract

The visual perception of 3D depth is underpinned by the brain’s ability to combine signals from the left and right eyes to produce a neural representation of binocular disparity for perception and behavior. Electrophysiological studies of binocular disparity over the past two decades have investigated the computational role of neurons in area V1 for binocular combination, while more recent neuroimaging investigations have focused on identifying specific roles for different extrastriate visual areas in depth perception. Here we investigate the population receptive field properties of neural responses to binocular information in striate and extrastriate cortical visual areas using ultra-high field fMRI. We measured BOLD fMRI responses while participants viewed retinotopic-mapping stimuli defined by different visual properties: contrast, luminance, motion, correlated and anti-correlated stereoscopic disparity. By fitting each condition with a population receptive field model, we compared quantitatively the size of the population receptive field for disparity-specific stimulation. We found larger population receptive fields for disparity compared with contrast and luminance in area V1, the first stage of binocular combination, which likely reflects the binocular integration zone, an interpretation supported by modelling of the binocular energy model. A similar pattern was found in region LOC, where it may reflect the role of disparity as a cue for 3D shape. These findings provide insight into the binocular receptive field properties underlying processing for human stereoscopic vision.

## 3 Introduction

Binocular stereopsis underlies our perceptual experience of stereoscopic depth and visual three-dimensional structure. Stereopsis is supported by a set of neural mechanisms for disparity selectivity and binocular integration that are distributed across multiple cortical regions in the human visual cortex (Backus et al. 2001; Bridge and Parker 2007; Preston et al. 2008; Ip et al. 2014; Goncalves et al. 2015; Li et al. 2019) and characterized by selective responses to specific stimulus features, such as absolute and relative disparity, surface curvature, slant, or separation in depth (Parker 2007).

To fully understand stereopsis, it is necessary to establish the relevant region of visual space over which binocular disparity is computed. We define this as the binocular integration zone, comprising the coincident retinal spaces of left and right eyes over which the two monocular inputs are pooled to form a unified binocular representation of disparity in cortical processing (Parker et al. 2016). The site of binocular combination can be localized to the primary visual cortex (V1) in the macaque monkey (Cumming and Parker 1999, 2000; Parker and Cumming 2001), with extrastriate areas performing computations relevant to binocular perception on an integrated representation of the binocular signal. Properties of the binocular integration zone could potentially be similar to the receptive field properties of responses driven by luminance contrast. Alternatively the binocular integration zone could differ in its spatial or temporal properties, revealing limits specific to disparity processing (Prince et al. 2002a; Nienborg et al. 2004, 2005; Anzai et al. 2011).

Neighboring binocular neurons display similar disparity selectivity, leading to clusters of cells encoding near or far disparities (Chen et al. 2008, 2017). This is compounded by the retinotopic organization of visual cortex, leading to regions preferentially responding to a particular binocular disparity, at a particular retinal location. This population-level organization makes disparity selectivity amenable to study with fMRI, a technique that samples cortical responses with a spatial resolution in the range of 1-2 millimeters. Neuroimaging studies of binocular disparity have characterized the spatial selectivity for binocular information across human visual cortex (Backus et al. 2001; Neri et al. 2004; Preston et al. 2008; Minini et al. 2010; Cottereau et al. 2011; Ip et al. 2014; Ban and Welchman 2015), within cortical areas (Nasr et al. 2016; Tootell and Nasr 2017), as well as the role they play in perceptual judgements of disparity (Backus et al. 2001; Goncalves et al. 2015; Bridge 2016). A previous study by Barendregt et al. (2015) found that a dichoptic bar stimulus presented in spatially offset positions in the two eyes led to larger population receptive fields in V1 compared to extrastriate regions. What remains unclear is how the characteristics of the binocular integration zone in V1 and extrastriate regions perform spatial integration when stimulated with pure binocular disparity, without any monocular cues compared to receptive fields driven by luminance and contrast.

We analyzed quantitatively the receptive field properties of binocularly driven receptive fields with population receptive field (pRF) methods. Using dynamic random dot stereograms in which sequential retinotopic positions are stimulated by changes in binocular disparity, we derived a pRF spatial model of fMRI signals that are specific to processing of a particular binocular disparity. The output of the pRF model summarizes the spatial extent of retinal locations over which a disparity signal increases cortical responses. This, in turn, is dictated by the properties of the neurons falling within the stimulated population, specifically the disparity selectivity and spatial selectivity of the receptive fields. As the maximum extent of the disparity-defined pRF is limited by the population-level binocular integration zone of binocular neurons in the sampled cortical space, we take the estimated pRF size as a valid estimate of the binocular integration zone. Sampling across multiple cortical visual areas, we show a pattern of larger pRFs for disparity-defined compared to checkerboard and luminance-defined stimuli in the

primary visual area V1 and also in the lateral occipital complex (LOC), supporting a distinct role for this extrastriate region in disparity processing. Model simulations reveal that the increase in estimated pRF size to disparity-defined stimuli in primary visual cortex is predicted by a standard binocular energy model (Ohzawa et al. 1990; Cumming and Parker 1997; Anzai et al. 1999).

## 4 Materials and methods

### 4.1 Participants

Eight healthy participants with normal or corrected-to-normal vision took part in the study (mean age 27.6 yr, age range 19-42, 6 female). They were screened for normal visual acuity (Snellen chart at 6 meters, <20/20 corrected) and stereoscopic vision (TNO test, <60 arcsec correct detection). This study received ethical approval from the University of Oxford Central University Research Ethics Committee (MS-IDREC-C1-2015-040) and was conducted in accordance with the Declaration of Helsinki (2013 revision). One participant was unable to successfully fuse the stereoscopic images, so they were not included in any of the analyses, leaving 7 participants included in the results.

### 4.2 Stimulus presentation

Visual stimuli were generated in MATLAB (v8.0, Mathworks Inc., Natick, MA, USA) using Psychtoolbox (v3.0, http://psychtoolbox.org) and displayed through a LCD projector (LC-XL 100, Eiki Industrial Company, Japan) via a back-projection screen situated inside the bore of the MRI scanner (peak luminance = 552 cd/m^2^). All stimuli were viewed through red and green anaglyph filters (Wratten 2 Optical Filters #29 and #61, Eastman Kodak, Rochester, NY, USA), both to provide stereoscopic display in the case of disparity-containing stimuli, and to ensure equal luminance attenuation across conditions. Luminance crosstalk, defined as the percentage of unintended signal to intended signal, was measured for the red and green filters at 0.16% and 0.82%, respectively. The red filter was always placed over the left eye.

Stimuli were arranged across five conditions (Figure 1): checkerboard, correlated disparity, motion, luminance, and anti-correlated disparity conditions. All stimuli were presented within the confines of a ‘wedge’ or ‘ring’ aperture, similar to that used in a standard retinotopic mapping design (Engel et al. 1994; Sereno et al. 1995). Four configurations were used: two types of wedge, rotating either clockwise or counter-clockwise, and two types of ring, either expanding or contracting. In the following section, we refer to the stimulus content within the aperture as the foreground and stimulus content outside the aperture as the background. The checkerboard condition (Figure 1A) consisted of a foreground of radial contrast-reversing (2Hz) checkerboard (contrast = 100%), while the background was set to 50% luminance, matched to the mean of the foreground stimulus. Stimuli were viewed binocularly through the red-green anaglyph filters.

**Figure 1.**
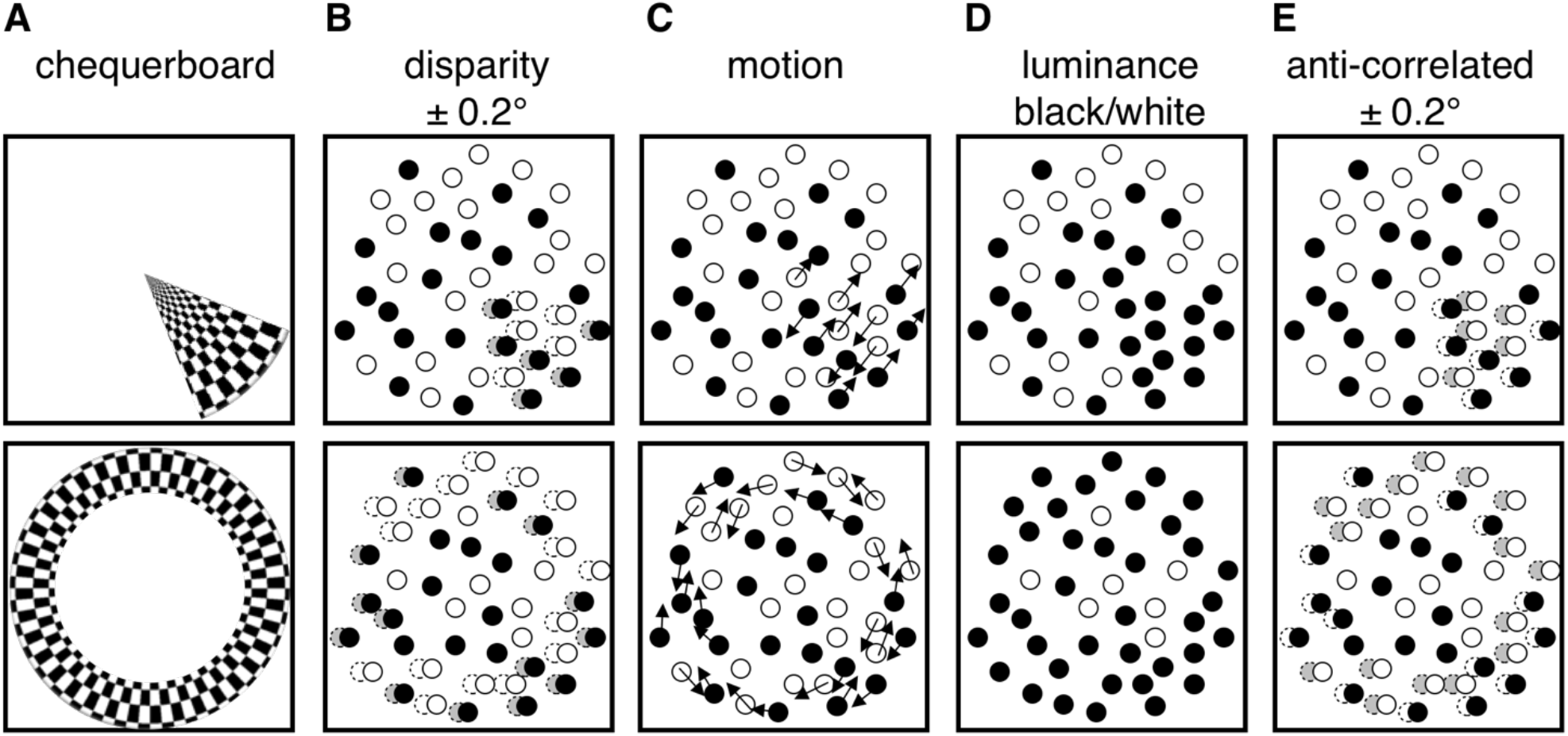
Experimental stimulus cartoon. Stimuli consisted of a contrast-reversing checkerboard (A) or dynamic random dot field (B-E) presented in a dynamically varied aperture of a wedge (upper row) or a ring (lower row). (A) Radial checkerboard reversing in contrast at 2Hz; (B) Disparity with random dot stimuli, dots inside aperture were binocularly correlated and changed in disparity (±0.2° from fixation); (C) Motion with clockwise and counter-clockwise components (dot speed = 7°/s); (D) Luminance reversal between fully black and fully white; (E) Anti-correlated binocular disparity (±0.2° from fixation).

The correlated disparity stimulus (Figure 1B), consisted of a dynamically changing array of randomly placed dots, half of them white and half black on a grey background. Foreground dots were fully correlated in position between left and right eyes and modulated in binocular disparity. Foreground dots were presented at either +0.2° or -0.2° disparity, corresponding to near and far positions relative to the fixation plane, and swapping every 1.45s. Background dots were randomly placed in left and right eyes and were therefore uncorrelated binocularly. Both foreground and background contained black and white dots (50% each), and dots refreshed at a frequency of 60 Hz.

The motion stimulus (Figure 1C) consisted of a dynamic random dot array, with dot positions fully correlated binocularly. Background dots were static, while foreground dots moved in either clockwise or counterclockwise motion (50% of dots in each direction) at 7°/s. To ensure dot motion was visible, dots were refreshed at a rate of 0.33 Hz, slower than the 60 Hz refresh rate for the correlated disparity stimulus. Both foreground and background contained black and white dots (50% each).

The luminance stimulus (Figure 1D) consisted of a dynamic random dot array, with dot positions fully correlated binocularly, refreshing at a rate of 60 Hz, and containing both black and white dots (50% each). Foreground dots were either 100% black or 100% white, with the luminance of foreground dots reversing at a rate of 0.69 Hz.

The anti-correlated disparity stimulus (Figure 1E) consisted of a dynamic random dot array, which was identical in layout to the correlated disparity stimulus (Figure 1B), except for the arrangement of foreground dot colors. Dots falling inside the aperture always had opposite contrasts in left and right eyes, so that they were presented in matching positions but displayed as either white in the left eye and black in the right eye, or the opposite. This manipulation negates the sensory percept of depth, while retaining binocular disparity information that is registered by V1 neurons (Cumming and Parker 1997). All stimuli were viewed binocularly through the red-green filters, with dynamic random dot arrays presented at a dot density of 40%, and a dot radius of 0.12° for 6 participants and 0.15° for 2 participants.

In addition to the main experimental stimuli, a single full-field radial checkerboard stimulus alternating with a grey background (2.5 s ON, 30 s OFF) was used to estimate the hemodynamic response functions (HRF) of visual cortex individually for each participant.

In order to control participants’ attention, a fixation cross was present throughout stimulation, and participants were required to detect a change of color of this cross from black to red. The fixation cross was presented in a radial 0.5° cut-out for all stimuli, and therefore any reconstructed pRFs with eccentricities <0.5° were discarded, due to overlap with the fixation cross. The color change was brief (200 ms) and occurred pseudo-randomly 80-100 times during a single run. Participants responded to this vigilance task via an MRI-compatible button box, and responses were monitored to ensure participant alertness. The average percentage of events detected was 88% ±5% SEM.

In all cases, modulation was displayed first within a wedge-aperture rotating clockwise or counterclockwise around the apex of the wedge, and subsequently within a ring-aperture expanding from or contracting into the centre. Dot positions were randomly generated for every frame, with 50% black and 50% white, except for the luminance modulation stimulus (for which dots are illustrated in black for clarity). Periods of no modulation (24/168 volumes per run) with a blank grey screen were used to estimate baseline response.

### 4.3 MRI acquisition

MR images were acquired with an ultra-high field 7T MRI system (Siemens Healthcare, Germany) using a 32-channel head coil (Nova Medical, USA). Functional imaging during visual stimulation was conducted with a gradient echo echo-planar imaging sequence (TR = 2488 ms, TE = 27.8 ms, 64 slices, resolution = 1.2 mm isotropic) with in-plane acceleration using parallel imaging (GRAPPA factor = 2) (Griswold et al. 2002) and through-slice acceleration using multiband imaging (MB factor = 2) (Moeller et al. 2010). Four runs were acquired for each stimulus condition, totaling 672 volumes per condition. The order of conditions and aperture order was randomized within and across sessions. For HRF estimation, three runs were acquired per participant, total 234 volumes. B_0_ field maps were acquired in-plane in each run to correct distortions due to field inhomogeneity. (TR = 620 ms, TE_1/2_ = 4.08 / 5.1 ms, resolution = 2 mm isotropic). A T1-weighted (T1w) whole-brain anatomical image was acquired to reconstruct the cortical surface and anatomically localize functional data (MP-RAGE, TR = 2200 ms, TE = 2.82 ms, TI = 1050 ms, flip angle = 7°, slices = 176, resolution = 1 mm isotropic).

### 4.4 MRI pre-processing

Functional images for each participant were pre-processed with FSL (FMRIB Software Library v5.0.8; http://www.fmrib.ox.ac.uk/fsl). EPI images were corrected for distortions caused by magnetic field inhomogeneities using FUGUE (Jenkinson et al. 2012), image portions showing brain tissue were isolated, and corrected for participant motion by linear realignment to the middle time point of each run. Low-frequency fluctuations were removed using a high-pass filter with cut-off at 0.02 Hz. Each run was then registered to the subject-specific T1w structural image using boundary-based registration (Greve and Fischl 2009).

Cortical surfaces were reconstructed from T1w structural images with FreeSurfer (v5.3.0, http://www.freesurfer.net). Volumes underwent automated segmentation to generate grey and white matter boundaries, and the grey matter surface reconstructed to create a two-dimensional representation of the cortical surface.

### 4.5 pRF analysis

fMRI data were analyzed using a Gaussian population receptive field (pRF) model (Dumoulin and Wandell 2008; Wandell and Winawer 2015). The analysis software was implemented in MATLAB and is described detail in (Alvarez et al. 2015). In brief, the participant-specific hemodynamic response function (HRF) was estimated by averaging 18 trials over the occipital lobes during full field checkerboard stimulation and fitting a double gamma function (Friston et al. 1995). Model predictions were constructed by combining the *a priori* position of the stimulus aperture at each MRI volume acquired and a radially symmetric two-dimensional Gaussian pRF. Predictions were then convolved with the participant-specific HRF and compared to the observed signal in a two-stage procedure. First, the spatially smoothed (full width half maximum = 5 mm, on spherical mesh) BOLD time courses were correlated with signal predictions generated by an exhaustive grid of combinations of the three pRF parameters (X coordinate, Y coordinate, σ size of pRF). The parameters resulting in the highest correlation at each vertex formed the starting point for the second stage, in which the original unsmoothed BOLD time courses were fitted using the Nelder-Mead algorithm for unconstrained nonlinear minimization (Lagarias et al. 1998) to identify parameter combinations for each vertex that maximize the variance explained by the model. Best-fitting model predictions yielded estimates of retinotopic location (X and Y coordinates) and pRF size (σ) for each vertex. Each condition (checkerboard, luminance, motion, correlated disparity, anti-correlated disparity) was fitted independently. Regions of interest were delineated for each participant based on polar angle and eccentricity estimates obtained in the checkerboard condition (see Figure 2).

**Figure 2.**
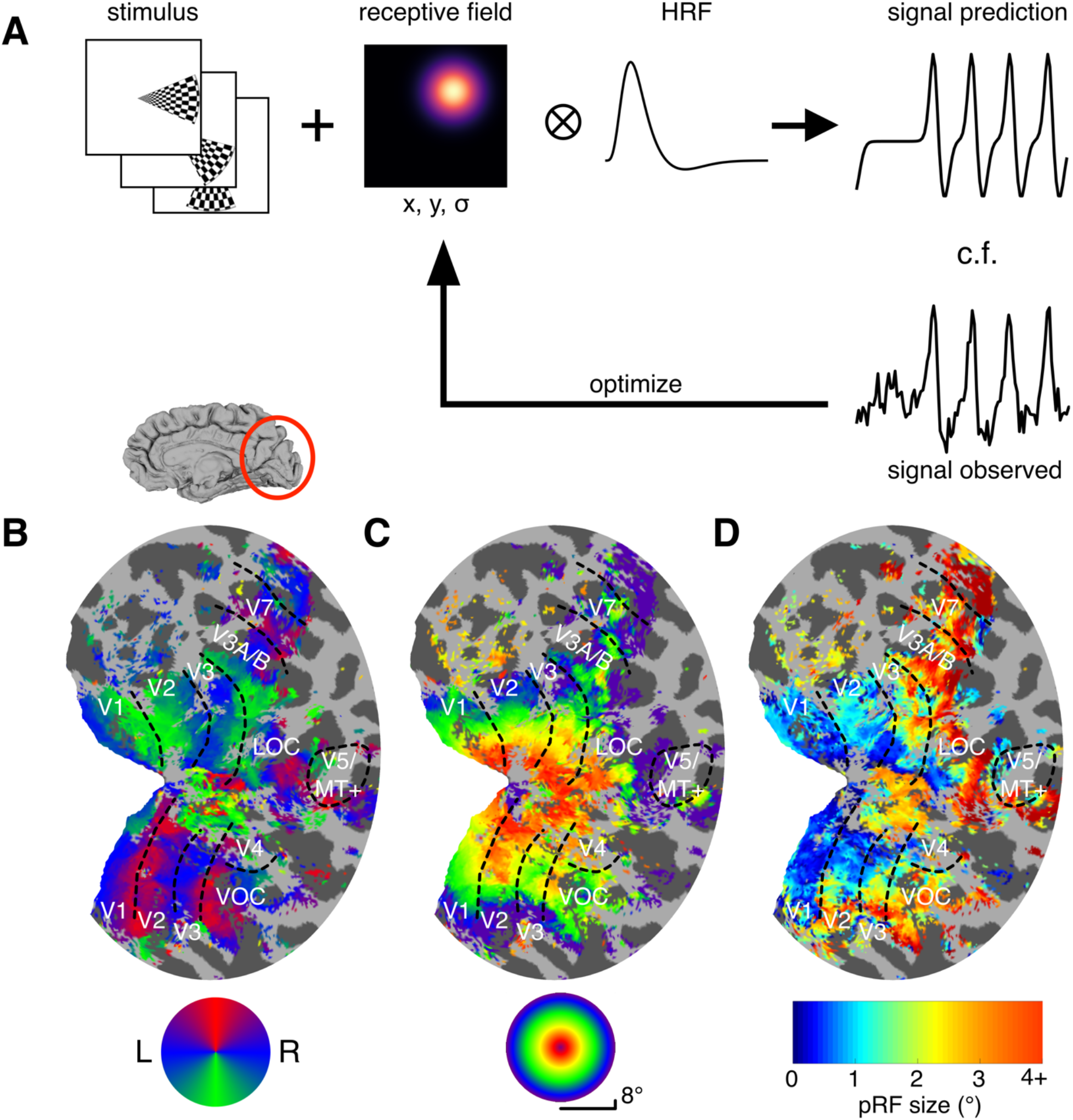
Cortical signals obtained under contrast-reversing checkerboard stimulation fitted with pRF model. (A) Model based on response to spatial and temporal sequence of visual stimulation of a Gaussian RF centered at location (X, Y) with spread (σ). Convolution with the participant-specific hemodynamic response function (HRF) gives a time-course prediction of the BOLD signal, which was in turn compared with the observed signal. Receptive field parameters (X, Y, σ) were then optimized iteratively to find the best-fitting pRF model for the data observed. (B) Polar angle delineation of visual areas; inflated right hemisphere for one participant (S1). Best-fitting model prediction shown. (C) visual field eccentricity of estimated pRFs. (D) pRF size across the visual cortex.

Model performance was assessed with the normalized correlation coefficient metric (CCnorm) (Schoppe et al. 2016). Each stimulus run was divided into two, with each half split considered an independent stimulus presentation. Signal reliability was used to normalize the correlation between pRF model prediction and empirically observed BOLD signals. Vertices were thresholded at CCnorm > 0.5, approximately equivalent to 50% of explainable variance explained by the pRF model.

### 4.6 Regions of interest

Regions of interest V1, V2, V3, V3A/B, V5/MT+, V7, V4, LOC and VOC were identified for each participant in each hemisphere tested. Since precise retinotopic boundaries could not be observed for all participants in some portions of visual cortex, a merged-region definition was adopted for areas LOC, VOC and V5/MT+. Specifically, the lateral occipital complex (LOC) encompassed retinotopic definitions of areas LO-1 and LO-2, the ventral occipital complex (VOC) encompassed areas VO-1 and VO-2 (Larsson and Heeger 2006; Wandell et al. 2007; Winawer and Witthoft 2015), and the region V5/MT+ encompassed the temporal-occipital areas TO-1 and TO-2 (Amano et al. 2009). Further, the region V5/MT+ was compared with an atlas definition of human occipital area 5 (hOc5), a cytoarchitectonic correlate of area V5/MT+, for anatomical agreement (Malikovic et al. 2006). This comparison showed a minimum of 50% overlap between vertices in retinotopically-defined V5/MT+ and the atlas-based cytoarchitectonic definition of hOc5 in all hemispheres tested (Mean overlap *=* 74%, *SD* = 12%, *N* = 14 hemispheres). The variability in alignment between structural and functional markers of V5/MT+ in human cortex has been noted before (Large et al. 2016).

### 4.7 Experimental design and statistical analysis

Differences in model performance between stimulation conditions were assessed in two ways. First, the distributions of CCnorm values were pairwise-compared between conditions with independent Kolmogorov-Smirnov tests. Note that p-values are unsuitable as measures of distribution similarity in the case of KS statistics (Vermeesch 2013), so only effect sizes are presented. Second, differences in mean CCnorm between conditions were assessed with a repeated measures ANOVA, introducing stimulation condition and region of interest as within-subject variables and participant identity as between-subject variable. Estimates of pRF size were also assessed for each visual area using a mixed effects model implemented in Prism (GraphPad Software, San Diego, California) with stimulus condition and eccentricity bin as within-subject variables and participant identity as between-subject variable. The anti-correlated disparity condition was not included in this analysis as there were too few vertices for which the pRF model could be successfully fit. This mixed-effects model was used rather than a repeated-measures ANOVA to account for vertices where no pRF model could be fit, hereafter labeled as missing values. Across all stimulus conditions (excepting anti-correlated disparity), visual areas, participants and eccentricities, 2.1% of values were missing. The disparity condition had the greatest number of missing values at 4.9%, but no individual visual area was missing more than 10% of values. Geisser-Greenhouse correction was applied where necessary and where random effects were zero, the term was removed and a simpler model fit used. Where there was a significant effect of condition in the main analysis, post-hoc mixed-effect models were conducted to assess the effect of specific condition pairs at each region of interest, with eccentricity bin introduced as a nuisance variable. All other statistical tests were implemented in MATLAB or SPSS (v24, IBM Corp., Armonk, NY, USA).

### 4.8 Binocular energy model

The response of binocular neurons in V1 to disparity information has been previously characterized as a set of canonical computations, formalized in the binocular energy model (Ohzawa et al. 1990; Cumming and Parker 1997; Anzai et al. 1999). In brief, the monocular inputs from left and right eyes arriving at a binocular simple V1 cell are each passed through a an eye-specific, weighting function approximated by a two-dimensional Gabor filter. Such filters are used in paired combination to capture the response to inputs to the left and right eyes, with horizontal displacement between the eye-specific filters conferring sensitivity to binocular disparity. A group of such filter pairs is set in quadrature, that is, with the spatial phase of the sinusoidal element of the Gabor functions offset by π/2 radians. Paired responses are summed, followed by a half-wave rectification and squaring nonlinearity. The summed output reflects the model binocular complex cell response (Ohzawa et al. 1990).

A population of 1,000 binocular complex neurons was simulated, with receptive fields positioned in the center of the visual field. A small normally distributed position offset (SD = 0.1°) was introduced to eliminate, by averaging, the spatial response of the population to detailed positions of the dots forming the randomly generated RDS patterns. The horizontal size of the Gabor profile, orthogonal to grating orientation, was manipulated to simulate receptive field size increase with eccentricity. Spatial frequency and disparity tuning were similarly manipulated to simulate the experimentally determined range of V1 receptive field properties found in recordings from macaque visual cortex (see below). The vertical size of the filter, parallel to grating orientation, was set to 1.5 times the horizontal size across eccentricity, also based on V1 recordings in the macaque monkey (Ringach et al. 2003). All filters were vertically oriented.

The stimuli delivered to the model receptive field consisted of (1) binocularly presented, contrast-reversing checkerboards (2) binocularly correlated dots in the aperture with binocularly uncorrelated dots in the background and (3) opposite polarity zero-disparity dots in the aperture and same polarity, zero-disparity dots in the background, just like the checkerboard, correlated disparity and luminance stimuli respectively viewed by participants. Stimuli were presented through a sweeping bar aperture in 100 steps, to create a timeseries of responses to the transient presence of contrast or disparity information. Random dots were binocularly correlated within the aperture at the programmed binocular disparity, but were uncorrelated in the background, while in the luminance condition dots were opposite polarity within the aperture and matched polarity in the background. For these random dot stimuli, 1,000 unique RDS frames were generated at each aperture step, and responses averaged together. Resulting responses were fitted with the Gaussian pRF model described in section 4.5. As estimation of the receptive field location is not of concern here, the location parameters were fixed *a priori* and only the receptive field size was estimated (Figure 3).

**Figure 3.**
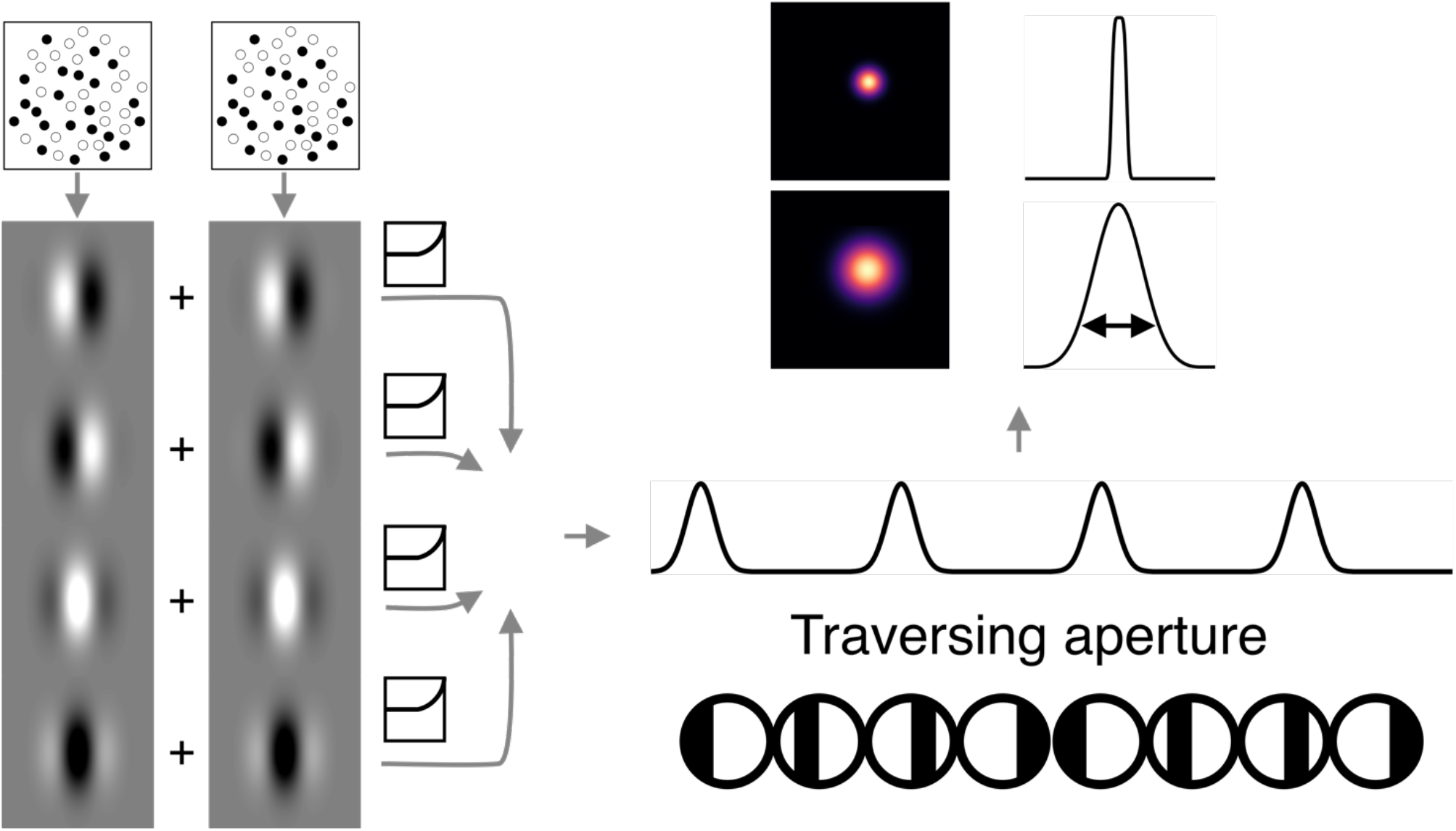
Binocular energy model implementation. Stimulus frames for the left and right eye were passed through a disparity-tuned Gabor filter bank, from which the monocular responses of each eye were linearly added, passed through a half-squaring non-linearity, and then pooled again by linear addition to produce the model complex cell response. The model cell responds

Three manipulations of model receptive field properties were conducted, in order to observe the effects on pRF model fits. First, the horizontal filter size was set to 15 different values between SD = 0.2° and SD = 3° to simulate RF size increase with eccentricity. The spatial frequency of the sinusoidal component of the Gabor was fixed to x0.5 the horizontal size, and all cells were set to be tuned to the stimulus disparity (checkerboard and luminance = 0°, disparity stimulus = 0.2°). Second, the same filter size points were sampled, while allowing spatial frequency of the Gabor filter to vary between x0.5 and x3.5 horizontal size, reflecting the variability in spatial tuning of V1 cells, while constrained by the size-disparity correlations observed in macaque V1 (Prince et al. 2002b). Third, both spatial frequency and disparity tuning were allowed to vary, with the latter allowing horizontal position of filters for left and right eyes to vary by SD ±0.25°. This final manipulation most closely resembles the distribution of receptive field properties reported for V1 cells in electrophysiological studies in macaque visual cortex. These simulations were designed to directly compare model responses generated by disparity-defined stimuli with checkerboard and luminance for the specific case when all model units are tuned to the stimulus disparity. A more comprehensive model would include units tuned to many different disparities, but this is beyond the scope of the current implementation.

periodically as the aperture defined by binocular correlation passes across its receptive field. The response timeseries of the population of RFs was fitted with the pRF model described. In the presence of a disparity-defined aperture, the width of the response is directly related to the area over which binocular information is integrated.

## 5 Results

### 5.1 Disparity responses are widespread across visual cortex

All visual cortical areas and regions of interest gave significant responses to binocular disparity stimulation as well as contrast stimulation. When considering the distributions of CCnorm, negligible effect sizes were detected when comparing the correlated disparity condition with the checkerboard (Kolmogorov-Smirnov distance, *KS* = 0.12, *D* = 10^−4^), motion (*KS* = 0.07, *D* = 10^−4^), luminance (*KS* = 0.07, *D* = 10^−4^) or anti-correlated conditions *(KS* = 0.15, *D* = 10^−4^). Variability in mean CCnorm was assessed with a repeated measure ANOVA, revealing a significant effect of condition (*F*(5,1) = 64.56, *p* = 10^−3^) and visual area (F(5,1) = 57.05, *p* = 10^−3^). Post-hoc *t-*tests showed all conditions outperformed the anti-correlated disparity condition (all comparisons *p* < 0.05), with no significant differences between the remaining conditions.

Figure 4 shows the pRF size averaged across all participants for each type of visual stimulation. The spatial distribution of pRF size estimates follows the expected pattern of small pRFs in areas representing the central visual field and larger in the periphery.

**Figure 4.**
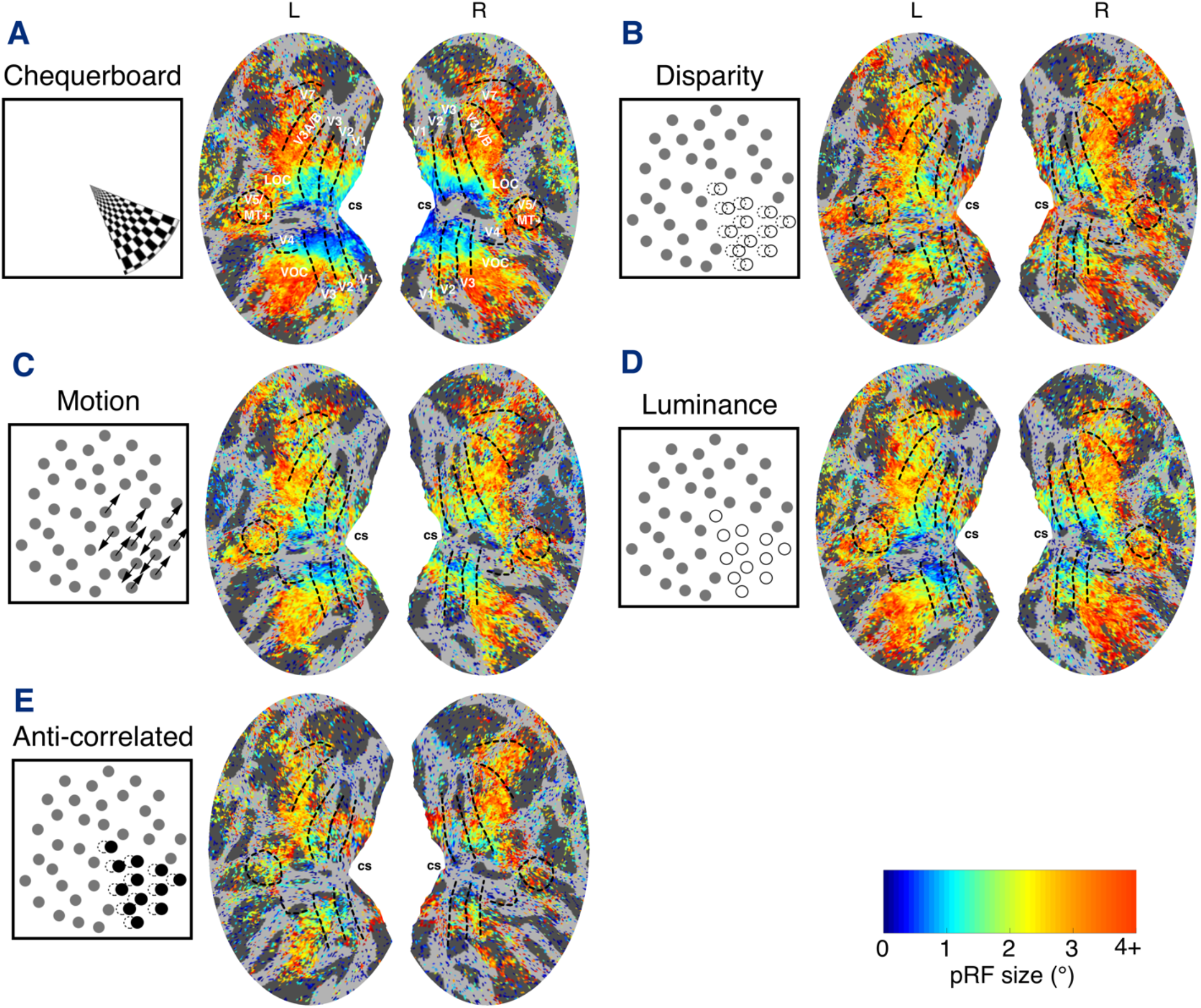
Group average estimates (*N*=7) of pRF size obtained under (A) checkerboard, (B) correlated disparity, (C) motion, (D) luminance and (E) anti-correlated disparity stimulation. pRF size estimates displayed on the normalized and flattened cortical surface, with cortical visual areas demarcated and calcarine sulcus (cs) labelled for reference. Vertices thresholded at CCnorm > 0.5.

### 5.2 pRF size for disparity varies systematically across the visual hierarchy

Estimates of pRF size obtained under the disparity condition may capture the binocular integration zone (Parker et al. 2016) over which monocular signals are combined, and consequently reflect the role that cortical areas play in the integration mechanism of binocular stereopsis. Examining the spatial distribution of pRF size estimates across the visual cortex reveals systematic variation: in particular, there are locations where pRF size estimates differ between the disparity and other conditions. We observed larger pRFs for correlated disparity in the calcarine sulcus, close to representation of the horizontal meridian, when compared to all other control conditions (Figure 4).

Binned estimates of pRF size for disparity across eccentricity are shown in Figure 5. Differences in binned values of pRF size between the correlated disparity condition and other conditions were assessed with a full factorial ANOVA model, introducing eccentricity and region of interest as independent variables. Anti-correlated responses were omitted from this comparison, owing to the low number of vertices successfully fitted by the pRF model under that condition. There was a significant interaction between condition and region of interest (*F* = 43.45, *df* = 14, 98, *p* = 0.001). In order to assess the effect of stimulus condition on a region-by-region basis, we conducted a series of linear mixed models, which are presented in the following section.

**Figure 5.**
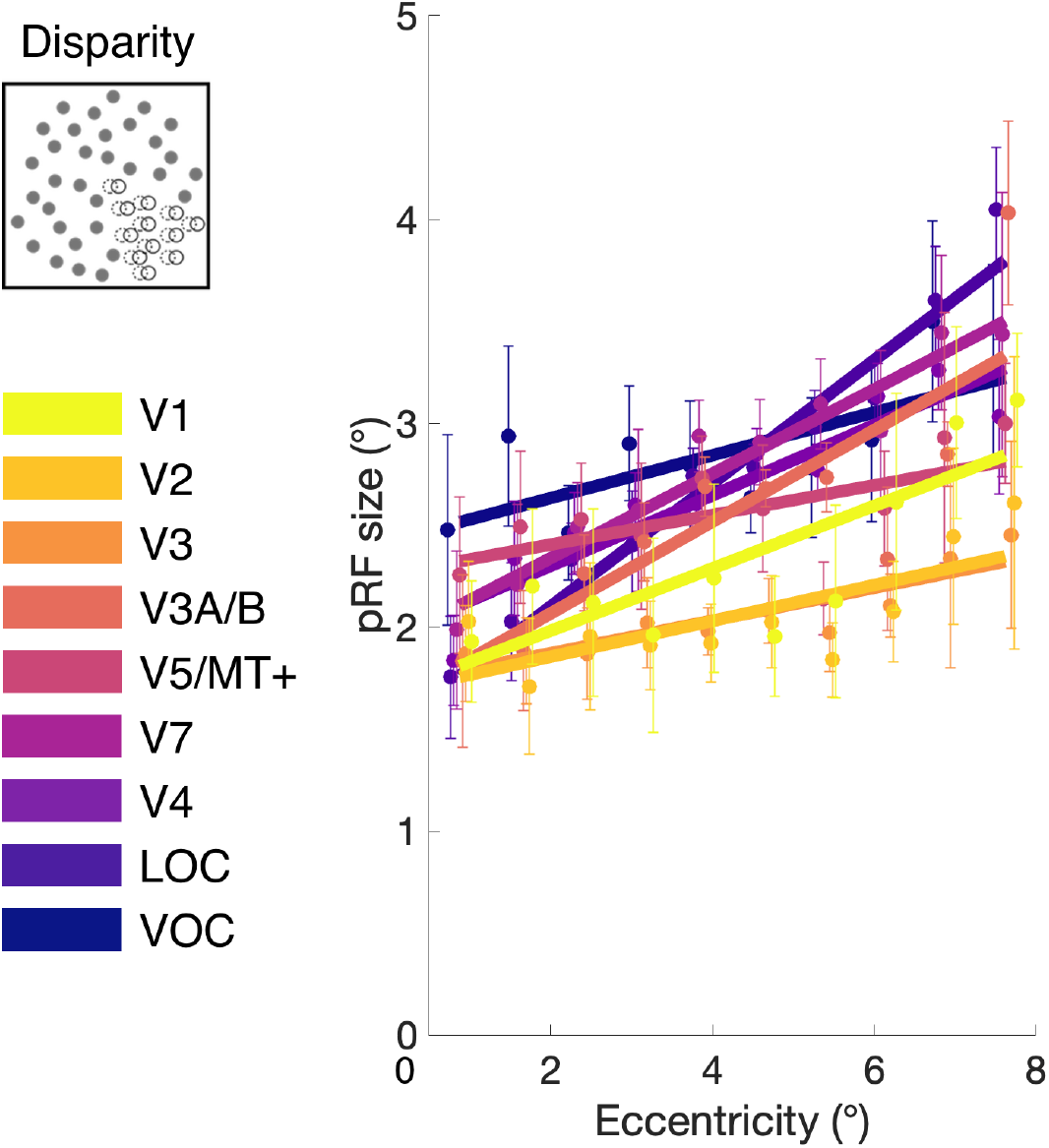
pRF size at different visual field eccentricities under disparity stimulation in the visual areas of interest. Estimates of stereoscopic pRF size were binned in 0.75° steps, with error bars indicating group SEM (*N*=7). pRF size increases both with eccentricity, and through the visual hierarchy. Vertices thresholded at CCnorm > 0.5. Small shift in eccentricity bin positions added to display differences between visual areas.

### 5.3 pRF size for disparity differs from non-disparity pRFs in V1

#### 5.3.1 Early visual regions: V1, V2 and V3

Figure 6 shows a summary of the pRF sizes at each eccentricity and for the different stimulus conditions across the early visual areas. In V1, a mixed effects model showed a significant effect of condition (F(2.1, 119.3) = 5; p = 0.007), although there was no effect of eccentricity (F(9, 60) = 1.9; p = 0.06) nor a significant interaction (F(27, 174) = 0.9; p = 0.62). Post-hoc paired comparisons showed that mean pRF size across all eccentricities for disparity (2.4°) were significantly greater than checkerboard (1.8°; F(1, 116) = 15; p = 0.0002) and luminance (2.0°; F(1,56) = 5.3; p = 0.02). Disparity pRFs did not differ in size from those defined by motion (2.1°; F(1,114) = 1.3; p = 0.26).

**Figure 6.**
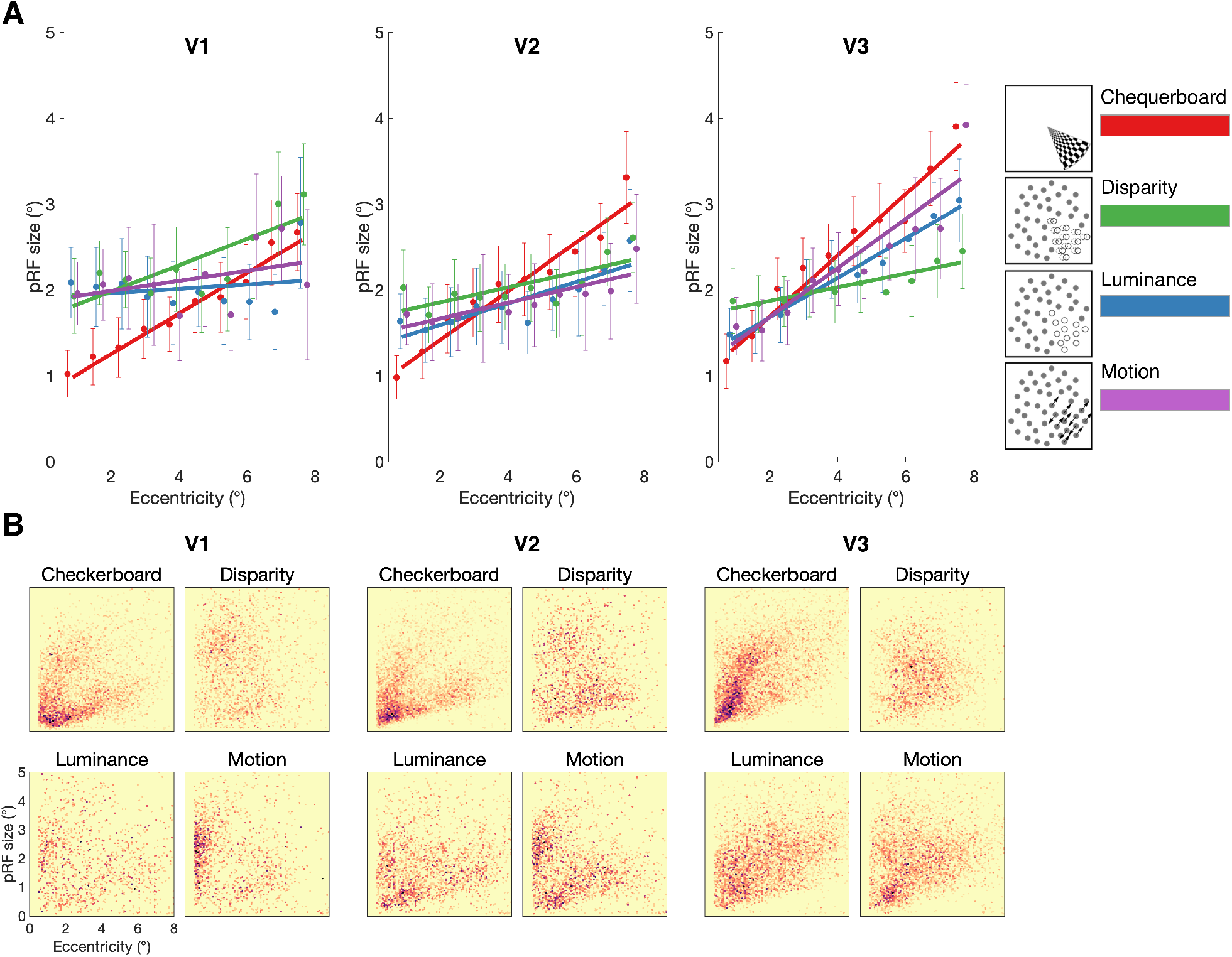
(A) pRF size estimates across visual field eccentricity for checkerboard, disparity, luminance, and motion stimuli, across visual areas V1, V2 and V3.pRF size values were binned in 0.75° steps and fitted with a linear regression model. Error bars indicate mean standard errors across participants. (B) Distribution of pRF size against visual field eccentricity across cortical surface points in all (N=7) participants tested. 2D binned histogram with 100×100 equal size bins. Darker colors indicate higher density of significant vertices.

In comparison, in V2, there was a significant effect of eccentricity on pRF size (F(9,60) = 3.8; p = 0.0008), but no difference according to stimulus condition (F(2.6,147.9) = 2.2; p = 0.11) or interaction (F(27,174) = 0.75; p = 0.81). Finally, in V3 there was a significant effect of eccentricity on pRF size (F(9,60) = 26.9; p < 0.0001), and stimulus condition (F(2.3, 136.6) = 5.5; p = 0.003) but no significant interaction (F(27,177) = 1.5; p = 0.06). However, while pRFs with the disparity stimulus (2.1°) were significantly smaller than with the checkerboard (2.5°; F(1, 117) = 16.2; p = 0.0001), they did not differ from either luminance (2.2°; F(1, 117) = 0.8; p = 0.37) or motion (2.3°; F(1, 57) = 3.1; p = 0.08)

Thus, it appears that only in V1 are pRF sizes greater for disparity than both checkerboard and luminance-defined stimuli. It is also the case in V1, and to a lesser extent in V2 that the checkerboard stimulus resulted in smaller pRF sizes at low eccentricities compared to the stimuli defined by dots. This effect may have been driven by spatial integration effects since the aperture boundaries formed by dot-defined stimuli require spatial integration over a larger region of the visual field compared to the stimuli with clear contrast-defined borders, such as the checkerboard stimulus.

#### 5.3.2 Dorsal visual regions: V3A/B, V5/MT+ and V7

The pRFs measured with the checkerboard stimuli were larger than those with the dot-defined stimuli across all dorsal visual areas (Figure 7). As evident from the graphs, there was a highly significant effect of eccentricity in all dorsal areas (V3A/B: F(9, 60) = 32.3; p < 0.0001; V5/hMT+: F(9,235) = 19.6; p < 0.0001; V7: F(9,236) = 38.6; p < 0.0001). Similarly, all areas showed a significant effect of stimulus type (V3A/B: F(2.3,136.3) = 11.1; p < 0.0001; V5/hMT+: F(2.0,158.0) = 4.4; p = 0.01; V7: F(2.7,214.5) = 16.3; p < 0.0001). However, the disparity-defined pRF size only differed from the pRF sizes defined using the checkerboard in V3A/B (F(1, 119) = 12.6; p = 0.0006) and V7 (F(1, 117) = 13.9; p = 0.0003). This suggests that any difference in these areas was more likely related to the use of dot-defined stimuli rather than disparity *per se*.

**Figure 7.**
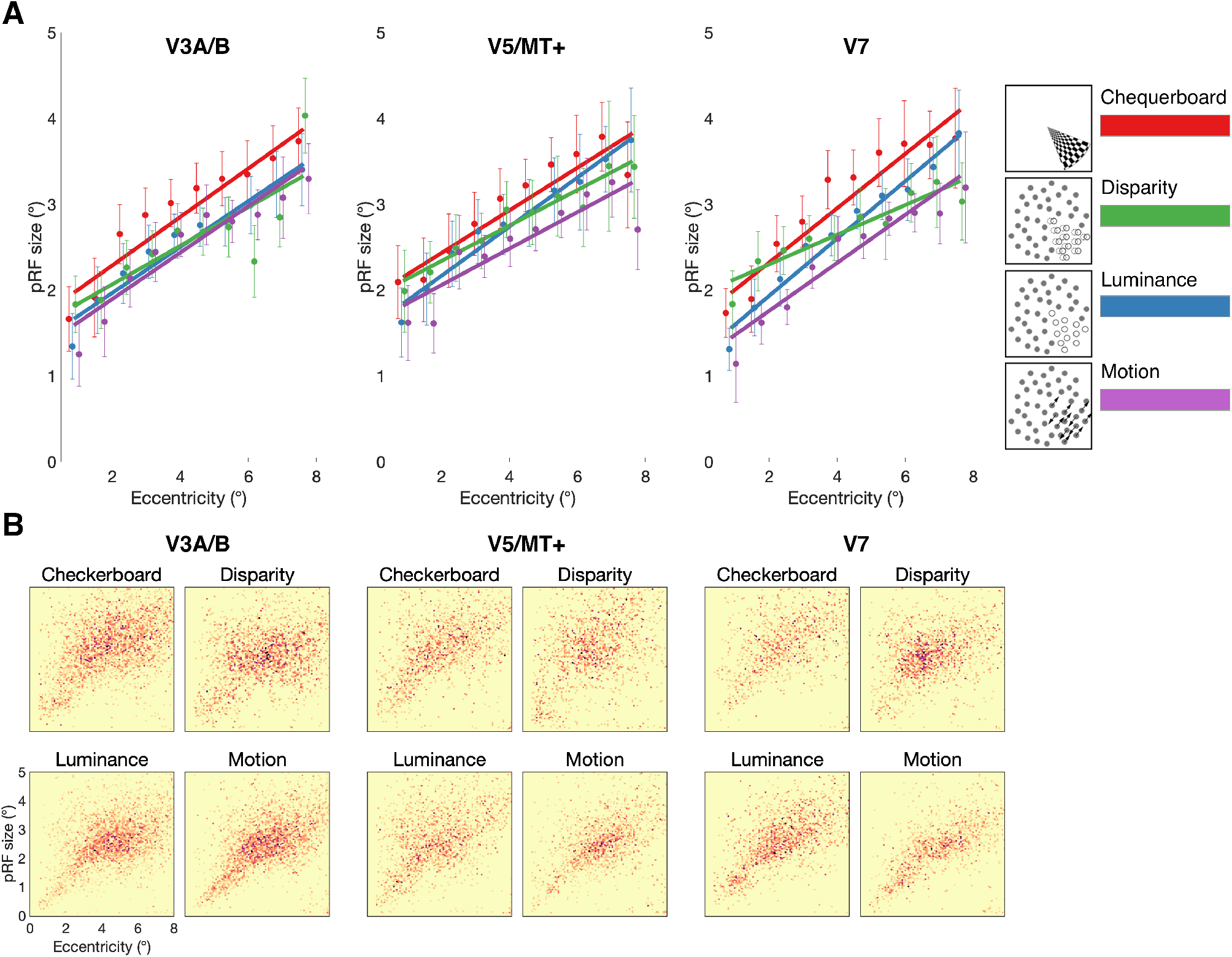
(A) pRF size estimates across visual field eccentricity for checkerboard, disparity, luminance, and motion stimuli, across visual areas V3A/B, V5/MT+, V7. pRF size values were binned in 0.75° steps and fitted with a linear regression model. Error bars indicate mean standard errors across participants. (B) Distribution of pRF size against visual field eccentricity across cortical surface points in all (N=7) participants tested. 2D binned histogram with 100×100 equal size bins. Darker colors indicate higher density of significant vertices.

#### 5.3.3 Ventral visual regions: V4, LOC and VOC

Figure 8 shows that in the ventral visual areas, like the dorsal regions, pRF size increased with eccentricity (V4: F(9, 230) = 16.6; p < 0.0001; VOC: F(9,60) = 5; p < 0.0001; LOC: F(9,60) = 50.3; p < 0.0001). There was a significant effect of stimulus type in both V4 (F(2.5, 193.0) = 10.8; p < 0.0001) and LOC (F(2.4, 137.4) = 5.4; p < 0.003) and V4 also showed a significant interaction between eccentricity and stimulus type (F(27, 230) = 2.4; p < 0.0002).

**Figure 8.**
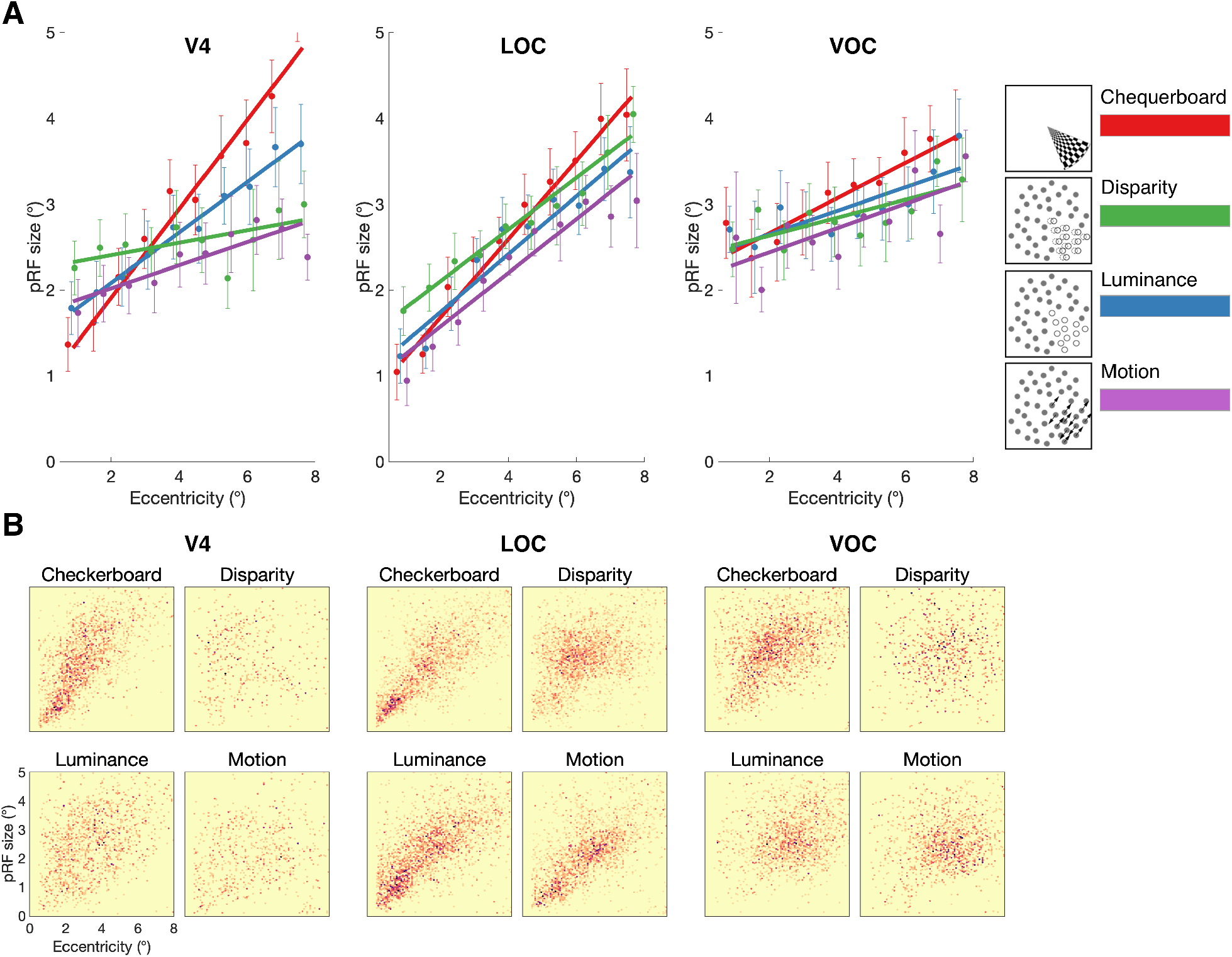
(A) pRF size estimates across visual field eccentricity for checkerboard, disparity, luminance, and motion stimuli, across visual areas V4, LOC and VOC. pRF size values were binned in 0.75° steps and fitted with a linear regression model. Error bars indicate SEM. (B) Distribution of pRF size against visual field eccentricity across cortical surface points in all (N=7) participants tested. 2D binned histogram with 100×100 equal size bins. Darker colors indicate higher density of significant vertices.

In V4, while there was a significant difference in pRF size when defined by disparity (2.6°) and checkerboard (3.1°; F(1, 112) = 12.5; p = 0.0006), disparity did not differ from either luminance (2.7°; F(1, 113) = 1.9) or motion (2.4°; F(1, 111) = 1.8). In contrast, pRF size in LOC was greater when defined by disparity (2.8°) compared to both luminance (2.5°; F(1, 44) = 10.3; p = 0.002) and motion (2.5°; F(1, 54) = 9.7; p = 0.003). When compared to checkerboard (2.7°), there was no difference in mean pRF size (F(1, 115) = 0.7), but there was a significant interaction (F(9, 115) = 2.4; p = 0.01) reflecting the larger pRF sizes for disparity at lower eccentricities, and smaller at the highest eccentricities.

### 5.4 Binocular energy model predictions of V1 integration zone

The disparity-defined stimulus used in the fMRI experiment contained a single magnitude of disparity (modulating from +0.2° to –0.2°), operating under the assumption that the estimated binocular integration zone would reflect the sub-population of binocular neurons that are tuned to these disparities, irrespective of the cortical territory being examined. However, electrophysiological studies have shown that disparity tuning co-varies with receptive field size, and by extension, with eccentricity (Prince et al. 2002a). This size-disparity correlation means the size of the binocular integration zone is dependent on eccentricity, as well as sensitivity to disparity-defined stimuli.

The binocular energy model provides a parsimonious account of the responses of binocular cells in area V1 in the presence of binocular disparity information, building disparity sensitivity from the linear combination of monocular receptive fields (Ohzawa et al. 1990; Cumming and Parker 1997; Anzai et al. 1999). In this model, the monocular receptive field is defined as a Gabor filter, that is, the product of a sinusoidal grating and a Gaussian envelope given by

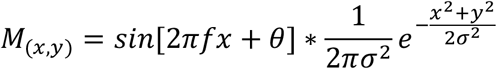

where *x* and *y* are point locations in 2D space, *f θ* is the grating phase, and σ is the standard deviation of the Gaussian envelope. In the presence of stimulus image *I*, the simple cell response is

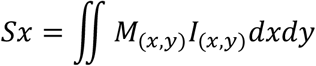

As the monocular receptive fields for the left and right eyes are independent, a position offset is introduced to generate sensitivity to binocular disparity. Summing and squaring the products of the monocular cells, create a linear-nonlinear ‘LN’ element

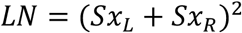

encoding disparity information at a particular phase and spatial frequency. A binocular complex cell is constructed by simply summing the LN elements of four pairs of monocular cells, set in phase-offset in quadrature

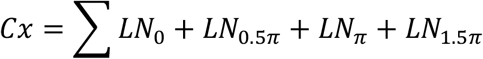

where two pairs of LN elements (θ = 0, π and θ =0.5π, 1.5 π) are anti-phase with each other. The response of the complex binocular cell can then be examined for the transient effect of stimulation; as matching retinal images overlap (or not) with the receptive fields, the response of the complex cell is modulated. In analogy with the fMRI task described above, this is equivalent to the stimulus aperture transiting across the model receptive field. Let us designate the aperture position *A*, for an arbitrary number of positions. In the case of a luminance-defined stimulus, the complex cell response is given by

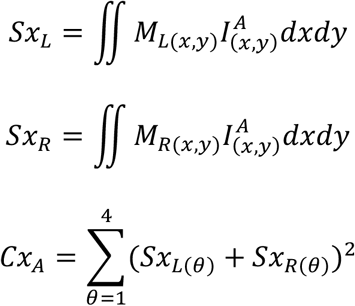

where the output of the complex cell is dependent on the overlap of the stimulus aperture *I*^*A*^ and the monocular receptive fields, *M*_*L*_ *and M*_*R*_. In the case of a non-disparity defined stimulus, the binocular receptive field reflects the simple sum of the monocular overlaps between the receptive fields. However, in the presence of stimulus disparity, a disparity-tuned complex cell pools information over an extended region of space, creating an expanded receptive field (Figure 9). In the presence of a transient aperture, this is translated into a wider window of response to disparity-defined stimuli. Thus, the relationship between the monocular receptive field width and the complex cell response to a transient stimulus is a function of disparity tuning.

**Figure 9.**
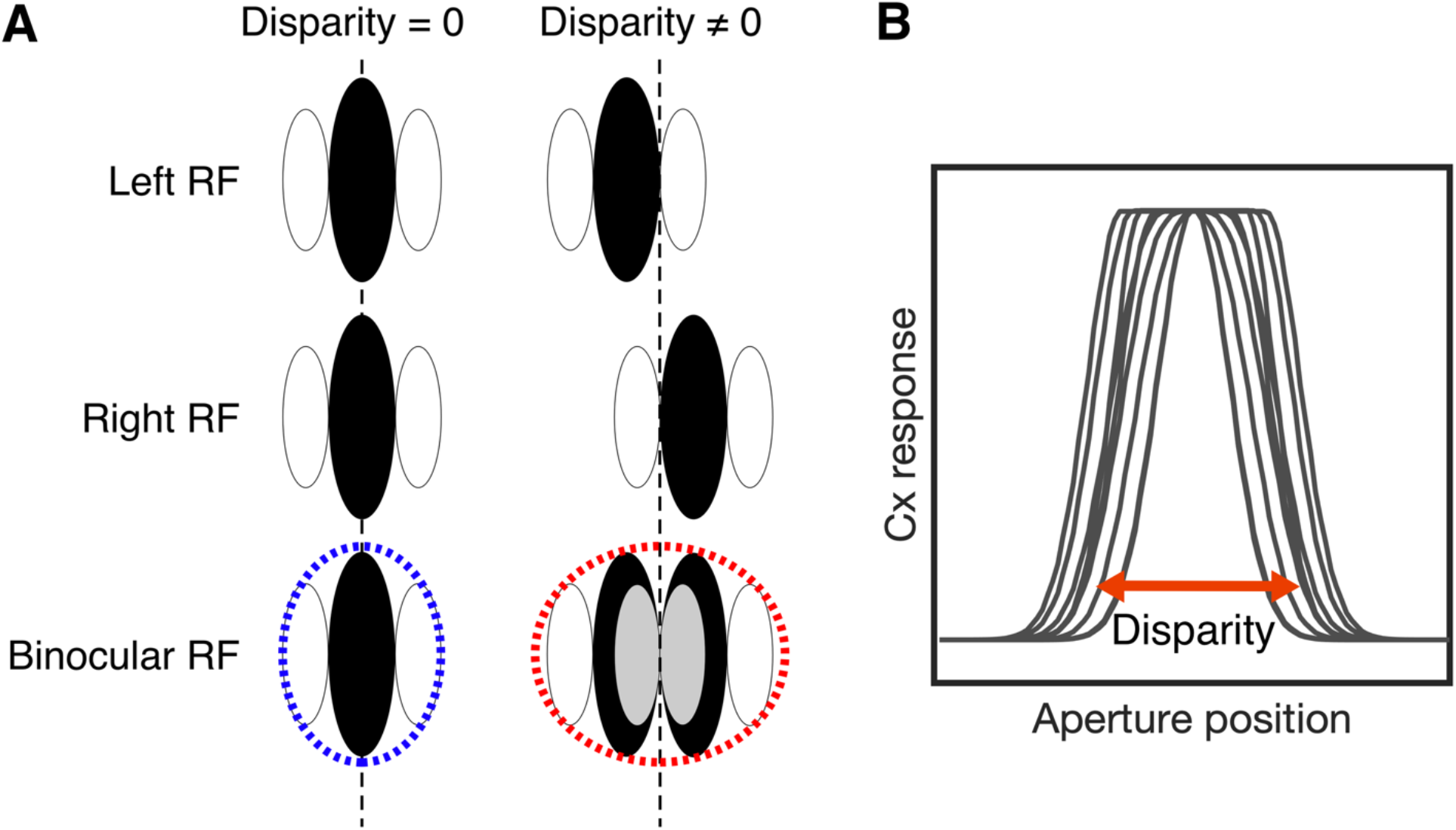
Binocular energy model construction in the presence of a transient disparity aperture. (A) The monocular receptive fields for the left and right eye are combined to form a binocular receptive field. Tuning to non-zero disparity increases the size of the binocular receptive field, proportional to the horizontal displacement between the monocular receptive fields. (B) In the presence of a disparity-defined transient aperture, the window of response from a complex binocular cell is determined by the magnitude of the stimulus disparity, the cell’s receptive field size and disparity tuning, i.e. the horizontal displacement between left and right monocular receptive fields. In the implementation presented here, both stimulus and model disparity are fixed at 0.2°

Implementing the binocular energy model described above, we examined the effect of modulating monocular receptive field sizes of a population of synthetic V1 neurons, and fitted the model responses with the pRF procedure, with results shown in Figure 10. A linear relationship between the model-defined receptive field size and the fitted binocular pRF size was observed, with responses to disparity-defined stimuli exhibiting larger pRFs compared to contrast-defined checkerboard and luminance-defined random dot stimuli. Using populations of model neurons with differing (i) receptive field size, (ii) spatial frequency, and (iii) disparity tuning produced a similar pattern of results similar discrepancies between stimulus content. These discrepancies qualitatively matched the pattern observed in the empirically estimated pRF sizes from BOLD data in area V1, which also displayed larger pRF sizes for disparity-defined stimuli when compared with contrast-and luminance-defined stimuli. Therefore, while size-disparity correlation limits the size of the binocular integration zone, these results support the view that the receptive field size, constrained by eccentricity, is the principal limiting factor on the size of the binocular integration zone.

**Figure 10.**
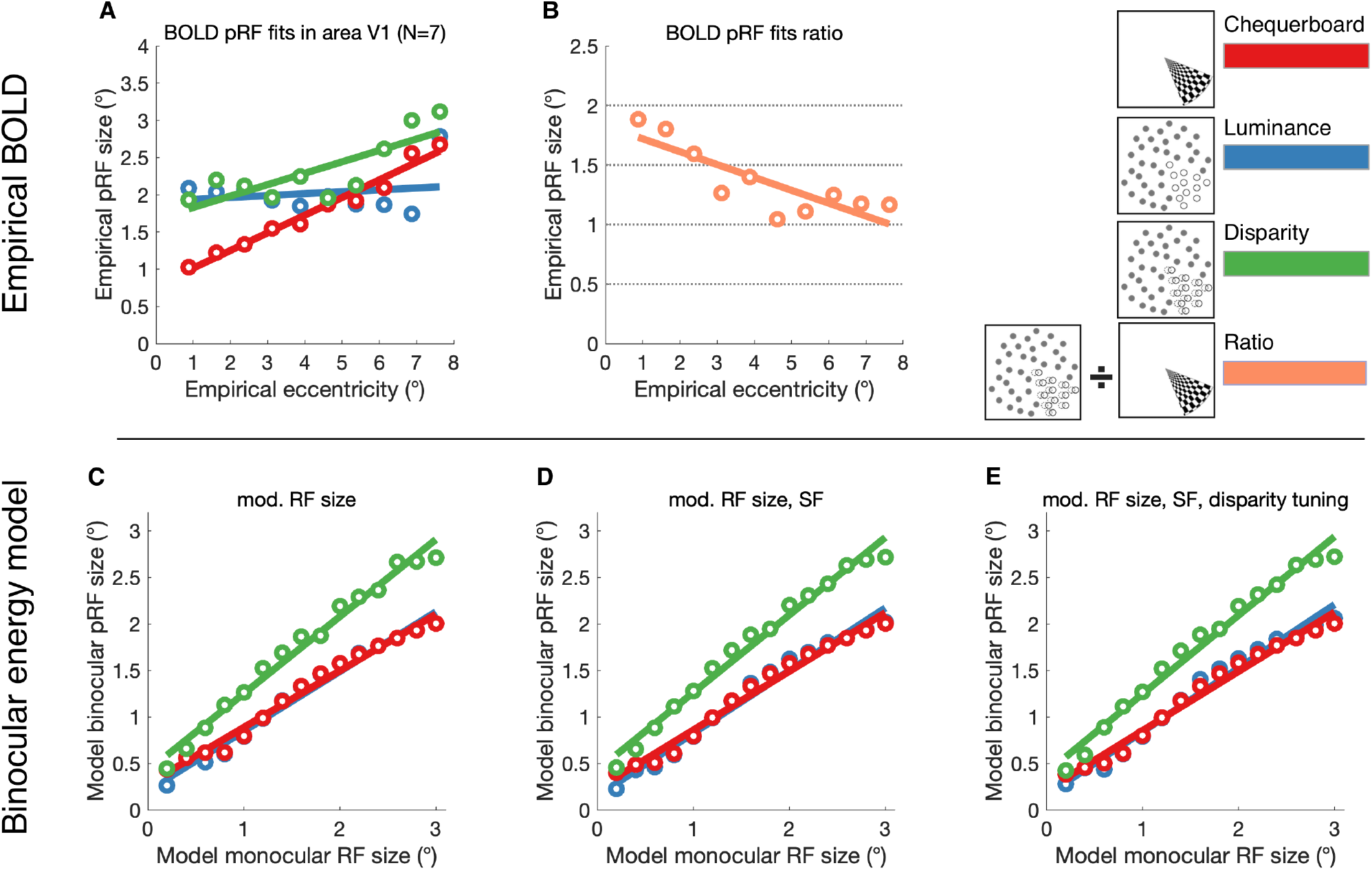
BOLD pRF fits compared to binocular energy model predictions. (A) Empirical BOLD pRF size increases with pRF eccentricity in area V1. pRFs measured under stimulation by disparity-defined RDS are larger than those measured contrast-defined checkerboard stimuli. Error bars omitted for clarity. (B) The ratio of pRF size for disparity over contrast is shown across eccentricities. (C-E) Binocular energy model predictions of simulated V1 cell populations show a similar pattern of pRF sizes, estimated by obtaining the model cell responses under disparity-, luminance- and contrast-defined stimulation and fitting the population-level responses with a pRF model. Varying model cell parameters across the model population in (C) receptive field (RF) size, (D) RF size and spatial frequency (SF) or (E) RF size, SF and disparity tuning, did not affect pRF size significantly under any type of stimulation.

## 6 Discussion

This study provides estimates of pRFs for binocular disparity across human cortical visual areas and compares them to estimates of pRF size for non-disparity defined stimuli. In particular, the derived pRFs obtained under correlated disparity stimulation are proposed to reflect the binocular integration zones of a given cortical site at the population level. Stimuli not defined by disparity, such as the luminance edges of the checkerboard, elicit responses across a wide variety of classical receptive fields, including both monocular and binocular RFs. By comparison, the stereoscopic RDS stimuli that define the wedge or ring aperture used to map the pRFs here only deliver aperture-related information encoded in binocular disparity. Therefore, where pRF estimates diverge between the disparity condition and pRFs estimated under luminance or contrast edges, differences should reflect the role of disparity-specific processing.

Our findings are consistent with previous fMRI evidence, which report widespread binocular disparity processing across visual cortex (Backus et al. 2001; Bridge and Parker 2007; Preston et al. 2008; Minini et al. 2010; Ip et al. 2014; Goncalves et al. 2015; Ban and Welchman 2015), and a specific role for area V1, as the site of binocular integration (Barendregt et al. 2015). An important point of interpretation for our study is that, unlike studies such as Barendregt et al., (2015), who compared binocular with monocular stimulation, the current study used binocular viewing in all tested conditions. Therefore, the stereoscopic stimuli used here probe the neuronal mechanisms that are responsible for the extraction of depth from binocular disparity. Second, our study makes direct comparisons of pRF size estimated in stereoscopic viewing-conditions for both disparity-and non disparity-defined stimuli, whereas the paper by Barendregt et al. (2015) compared overall quality of fits of the pRF model to the monocular and binocular stimulation conditions.

### 6.1 Estimates of the binocular integration zone in area V1

Our results demonstrate a discrepancy between pRFs estimated from disparity and non-disparity information in area V1, with larger receptive fields for disparity in agreement with the electrophysiological literature (Nienborg et al. 2004). This is consistent with the proposal that the binocular combination in disparity-specific neurons of V1 is a fundamental limiting stage in determining the size of the pRF (Cumming and Parker 1999, 2000; Parker and Cumming 2001). The lack of discrepancy of pRF size in areas V2 and V3 suggests little further combination of the retinal inputs in early extrastriate cortex, at least at levels detectable by population-level methods. In this regard, our findings are similar to those of Barendregt et al. (2015).

The relationship between the sizes of the non-disparity receptive field and the binocular integration zone in V1 is described by the general form of the binocular energy model (Banks et al. 2004; Nienborg et al. 2004). In this model, the ability of disparity-tuned V1 cells to detect changes in binocular disparity is limited by the width of the correlation window over which monocular signals are compared. If the window is too large, binocular matches become ambiguous; if the window is too small, the binocular image will not contain enough information to compute disparity (Banks et al. 2004; Nienborg et al. 2004). Notably, this constraint is independent of depth variation within the window, or limits imposed by optical effects, retinal sampling or stimulus construction (Tyler 1974; Schlesinger and Yeshurun 1998; Banks et al. 2004). As the correlation window is defined by the size and location of the paired monocular receptive fields, the latter impose the minimum area over which disparity information may be integrated. Indeed, the disparity energy model predicts a binocular integration zone whose effective receptive field is the half-squared product of the monocular receptive fields over which binocular cross-correlation takes place (Banks et al. 2004; Nienborg et al. 2004, 2005). This prediction is borne out in electrophysiological studies; for example, (Nienborg et al. 2004) showed that for disparity-tuned V1 neurons, the relationship between monocular receptive field size and the width of the correlation window corresponds to a half-squaring output nonlinearity and is approximately linear across eccentricities. Extrapolating this idea to neuronal population level, the binocular integration zone is predicted to display a half-square non-linearity in relation to other conditions, equivalent to a positive slope in the pRF size ratios between disparity and control conditions in Figure 6B.

### 6.2 Comparison of binocular energy model prediction and the empirical binocular integration zone

An explicit, but restricted implementation of the binocular energy model allowed us to assess the effects of monocular receptive field parameters on the conjugate signal of a model V1 cell population. By manipulating the model monocular receptive field size and fitting the mean population signal with a pRF model, we confirmed that pRF size for disparity is a linear function of monocular receptive field size. We also confirmed that disparity-defined stimuli resulted in larger pRFs compared to both contrast- and luminance-defined stimuli, reflecting the wider binocular integration zone necessary for integrating horizontal discrepancies in the monocular inputs, absent in the case of contrast information. Deriving the pRF from a population of model cells with different receptive field sizes, SF and disparity tuning produced a relationship between responses to disparity-defined and contrast-defined stimuli that was comparable to the empirically-estimated pRF sizes from BOLD data in area V1.

Nonetheless, there were several discrepancies between the empirical data and modeling results. Firstly, when comparing the pRF size for disparity with the checkerboard, in the model the difference between the two conditions appears to include with receptive field size, whereas the empirical data appears to converge. However, the model data indicate that at the largest sizes the curves being to converge. Secondly, the model pRF sizes for contrast and luminance stimuli show almost exactly the same pattern. In contrast, the empirical data for these two conditions varied considerably, with smaller pRFs at low eccentricities and larger pRFs at high eccentricities for the checkerboard. Why such a discrepancy exists is not clear, but may reflect salience of the stimulus, which is lower in the luminance condition.

Finally, as stated earlier, the current implementation of the energy model uses populations of units, but all have the same disparity tuning, which imposes limits the size of the pRF for the disparity-defined stimulus. As more units are incorporated that are tuned to different disparities, it is likely that the pRF size will increase, but this requires considerably more modeling that is beyond the scope of the current study.

### 6.3 pRF size is comparable for disparity and non-disparity input defined by random dots in dorsal visual areas

Dorsal regions V3A/B, V5/MT+ and V7 showed no significant difference in pRF size for disparity when compared to the dot-defined luminance and motion conditions. There was, however, a reduction in pRF size compared to the checkerboard stimulus. This is consistent with previous work indicating that pRF mapping with isolated dot-defined bar stimuli resulted in larger pRF sizes compared to stimuli presented with a contrasting surround, either opposing motion or motion noise (Hughes et al. 2019). Thus, given the lack of difference between disparity-defined stimulus and other dot-defined stimuli, our finding is consistent with the conclusion from that paper that the pRF size in dorsal regions may depend on stimulus salience. While this result is also consistent with significant involvement of dorsal visual areas in disparity processing, most notably V3A/B (Poggio et al. 1988; Adams and Zeki 2001; Neri et al. 2004; Minini et al. 2010; Ban and Welchman 2015), it does not indicate a special role for integration of disparity information across space.

### 6.4 Specialized processing for binocular disparity in lateral occipital cortex

In a similar fashion to the results observed in V1, we detected a pattern of larger pRFs for disparity compared to other conditions in area LOC, typically considered a later ‘upstream’ stage in visual cortical hierarchy processing (Grill-Spector et al. 2001). LOC is involved in the processing of 3D shape (Kourtzi and Kanwisher 2001; Kourtzi et al. 2003; Weigelt et al. 2007; Vernon et al. 2016), motion (Moutoussis et al. 2005; Krekelberg et al. 2005) and binocular depth (Chandrasekaran et al. 2006; Preston et al. 2008; Ban et al. 2012). While responsive to binocular disparity stimulation in isolation (Ip et al. 2014), LOC has been particularly associated with view-invariant representations of 3D shape which incorporate information about binocular depth (Welchman et al. 2005; Preston et al. 2009). The discrepancy between pRF sizes for disparity and other conditions may reflect the computational role for disparity information in LOC, not as the input to a binocular integration zone to generate a fused cyclopean representation, but instead as one component drawn upon to form view-invariant object representations. Preston et al., (2008) suggest that LOC represents depth position in a categorical manner, that is, as a coarse indicator of near vs. far position. As larger binocular disparities require larger receptive fields to capture the relevant retinal matches, it follows that coarseness in disparity tuning in LOC may be matched with a coarse spatial tuning in its pRFs. While the relationship between disparity tuning and receptive field size remains largely unknown in the human, in the macaque, electrophysiological studies have reported a multiplicative relationship between receptive field size and preferred disparity for V1 neurons (Prince et al. 2002b; Nienborg et al. 2004). Therefore, a coarse representation of both spatial and disparity tuning in LOC would be consistent with the tuning properties of disparity-selective cells.

An additional consideration is the source of disparity modulation. The dynamic random dot disparity stimulus presented here contains two sources of disparity information; absolute disparity within the aperture field, and relative disparity at the edge between the aperture and the zero-disparity background. Unlike area V1, which is exclusively selective to absolute disparity (Cumming and Parker 1999), either component may drive responses in LOC. While LOC responses can be attributed to relative disparity (Welchman et al. 2005; Chandrasekaran et al. 2006; Preston et al. 2008; Read et al. 2010; Bridge et al. 2013), a direct coding of absolute disparity is possible and consistent with the similarity in tuning properties with area V1.

### 6.5 fMRI estimates of binocular pRFs are in agreement with electrophysiological priors

This study presents the novel estimation of binocular receptive fields characteristics across human visual cortical areas, highlighting the discrepancies between disparity and non-disparity driven estimates of population-level receptive fields. While the estimates of pRF size for non-disparity modulated stimuli presented here are in broad agreement with previous fMRI studies (Wandell and Winawer 2015), no such baseline is available for disparity-defined pRFs. Furthermore, although direct comparisons to the electrophysiological literature may be informative, it is important to note the abstraction of these metrics from the behavior of single disparity-tuned cells. First, BOLD fMRI signals are measured from imaging voxels that contain many cells, both tuned and not tuned to disparity, which contribute to the observed signal. Second, imaged voxels encompass a large number of disparity-sensitive neurons that contain a variable distribution of spatial and depth preferences that are aggregated and averaged in the observed signal. Therefore, the BOLD signal reflects a population preference, which nevertheless reveals systematic variation in pRF size for disparity both within and across cortical visual regions.

Relating these findings to electrophysiology, we highlight two points. First, pRF size for disparity increased with eccentricity in all visual areas tested. Secondly, the scaling of pRF size with eccentricity under disparity stimulation is consistent with the view of a binocular integration zone that obeys both local physiological constrains imposed by its component receptive fields, and imposing a limit on resolvable disparity (Banks et al. 2004; Nienborg et al. 2004). Together, these observations reinforce the hypothesis that fMRI estimates of binocular receptive fields reflect the same mechanisms as those described in electrophysiological studies of disparity processing in animal models and provide the first characterization of the binocular integration zone in humans.

## 7 Declarations

### Funding

This work was supported by the Medical Research Council (MR/K014382/1), and The Royal Society (University Research Fellowship to HB). The Wellcome Centre for Integrative Neuroimaging is supported by core funding from the Wellcome Trust (203139/Z/16/Z).

### Conflict of interest

The authors declare no competing financial interests.

### Availability of data and material

The datasets generated during and/or analysed during the current study are available from the corresponding author on reasonable request.

### Code availability

The population receptive field modelling toolbox is available from https://github.com/samsrf/samsrf. The binocular energy modelling toolbox is available from https://github.com/IvanAlvarez/BinocularEnergyModel.

### Authors’ contributions

Conceptualization: IA, SAH, AJP, HB. Methodology: IA, SAH, AJP, HB; Formal analysis and investigation: IA. Writing -original draft preparation: IA; Writing -review and editing: IA, SAH, AJP, HB; Funding acquisition: AJP, HB.

### Ethics approval

This study received ethical approval from the University of Oxford Central University Research Ethics Committee (MS-IDREC-C1-2015-040) and was conducted in accordance with the Declaration of Helsinki (2013 revision).

### Consent to participate

Informed consent was obtained from all individual participants included in the study.

### Consent for publication

The authors affirm that human research participants provided informed consent for publication of results based on data collected in this study.

## Notes

### Competing Interest Statement

The authors have declared no competing interest.

